# Ventricular chamber-specific Pitx2 insufficiency leads to cardiac hypertrophy and arrhythmias

**DOI:** 10.1101/253062

**Authors:** Ana Chinchilla, Francisco J Esteban, Estefanía Lozano-Velasco, Francisco Hernandez-Torres, Jorge N Dominguez, Amelia E Arànega, Diego Franco

## Abstract

Genome-wide association studies (GWAS) have identified genetic risk variant adjacent to the homeobox transcription factor PITX2 in atrial fibrillation (AF) patients. Experimental studies demonstrated that Pitx2 insufficiency leads to cellular and molecular substrates that increased atrial arrhythmias susceptibility. Pitx2 expression is present not only in the atrial but also in the ventricular myocytes. This study aims to investigate if insufficiency of *Pitx2* in the developing and adult ventricular chambers increased susceptibility to ventricular arrhythmias. Conditional *Pitx2* loss-of-function ventricular chamber-specific (Mlc2v-Cre) mouse mutants were generated using Cre/loxP technology. *Pitx2* insufficiency in the ventricular myocardium leads interventricular septal thickening during cardiogenesis but else mice are viable until adulthood. Adult Mlc2vCre^+^Pitx2^-/-^ hearts display hypertrophic and dilated ventricular chambers. ECG recordings demonstrated that Mlc2vCre^+^Pitx2^-/-^ mice display increased QT and QRS intervals. Molecular analyses demonstrate that repolarization but not depolarization is severely impaired in these mutants. Microarrays analysis identified mRNAs and microRNAs altered in Pitx2 ventricular-specific mutants and provide evidences for miR-1 and miR-148 deregulation which in turn modulate Klf4 and distinct ion channel expression linked to cardiac hypertrophy and long QT-like defects. Our data demonstrate that Pitx2 insufficiency play leads to cellular and molecular ventricular remodeling which results in hypertrophic and dilated ventricular chambers and electrophysiological defects resembling long QT syndrome.

---

Pitx2 is a homeobox transcription factor that plays an essential role in the early left/right determination (1). *Pitx2* expression is confined to the left side of the embryo within the lateral plate mesoderm, and with further development *Pitx2* is maintained asymmetrically expressed in organ primordia such as the stomach and the heart (1-3). Thus, the homeobox transcription factor *Pitx2* is the only component in the early left-right signaling cascade that is expressed in the cardiac crescents and early heart tube (4).

The *Pitx2* gene generates three isoforms, *Pitx2a*, *Pitx2b* and *Pitx2c*, which are coded by alternative splicing (*Pitx2a* and *Pitx2b*) and by different promoter usage (*Pitx2c*) in mice (2). The generation of loss-of-function mouse mutants deleting all three *Pitx2* isoforms, i.e. *Pitx2abc* null, displays early embryonic lethality with severe cardiac malformations (5-7), demonstrating the importance of *Pitx2* during cardiogenesis. Ai et al. (8) have reported that lack of *Pitx2* in the second heart field results in similar developmental phenotype as the germline *Pitx2* mouse mutants. In addition, Ammirabile et al. (9) and Tessari et al. (10) has recently demonstrated that lack of *Pitx2* in distinct stages of myocyte determination results in distinct cardiac abnormalities during embryogenesis.

Gudbjartsson et al. (11) and Kääb et al. (12) have suggested a new role for PITX2 in the adult heart (see for a recent review Franco et al. (13)). GWAS have independently reported several risk variants on chromosome 4q25 that are strongly associated with AF in distinct human populations (14-20). Experimental evidences demonstrated that impaired Pitx2 expression in mice increased atrial arrhythmias susceptibility (21-25). Experimental AF similarly disrupts PITX2 expression in pigs (26) further demonstrating a pivotal role of Pitx2 in atrial electrophysiology. Pitx2 is not confined to the atrial chambers but expression is also present within the ventricular chambers (4, 9, 23). In this study we explored whether Pitx2 contributes to cardiac arrhythmias in the ventricular chambers. By generating Pitx2 ventricular-specific conditional mutant mice, we demonstrate that ventricular deletion of Pitx2 leads to developmental defects which are compatible with life but predispose to impaired electrophysiological function of the adult heart, reminiscent of long QT syndrome. Molecular analyses demonstrate that potassium, but not sodium, ion channel expression is impaired. Microarrays analysis identified mRNAs and microRNAs altered in Pitx2 ventricular-specific mutants providing evidences that Pitx2 modulates miR-1 and miR-148, which modulate Klf4 and distinct ion channels, thus contributing to both arrhythmogenic and hypertrophic features of Pitx2 ventricular-specific insufficiency.

## MATERIALS AND METHODS

### Transgenic mouse lines, breeding strategy and PCR screening

The Pitx2floxed and Mlc2v-Cre transgenic mouse lines have been previously described (7, 27). Generation and PCR screening of conditional ventricular (Mlc2v-Cre) mutant mice was performed as detailed in Supplementary data. Since conditionally-deleted homozygous mice were viable to adulthood, mice were bred into C57Bl/6J genetic background during 10-12 generations, and offspring was routinely screened for the presence of the Pitx2 floxed allele and the Cre recombinase sequence as previously reported (9, 10, 23). This investigation conform the *Guide for the Care and Use of Laboratory Animals* published by the US National Institutes of Health. The study was approved by the University of Jaen Bioethics Committee.

### Anatomical and histological analyses

Mice were sacrificed in all cases by cervical dislocation. Adult hearts were carefully dissected and briefly rinsed in Ringer’s solution and photographed. Samples processed for histochemistry and immunohistochemistry were fixed overnight in freshly made sterile 4% parafomaldehyde. Samples processed for RNA isolation were dissected (if applicable) and immediately snap-frozen in liquid nitrogen and stored at ‐80°C until used. Staged E13.5 and E16.5 embryos and were carefully dissected from uterus, briefly rinsed in sterile PBS and processed accordingly. Adult and embryonic samples processed for histochemistry and immunohistochemistry were dehydrated through graded ethanol steps and embedded in paraplast. Sections were cut at 10 μm and processed for hematoxylin and eosin, Mason´s trichrome and picrosirius according to standard procedures.

### Microarrays analysis

mRNA microarrays (CodeLink) analyses were performed in triplicates to profile mRNA signatures between E16.5 control (Mlc2vCre^-^Pitx2^fl/fl^) and homozygous conditional (Mlc2vCre^+^Pitx2^-/-^) embryos. Tissue samples were collected and pooled from isolated ventricular chambers (n=5) corresponding to E16.5 control (Mlc2vCre^-^Pitx2^fl/fl^) and homozygous conditional (Mlc2vCre^+^Pitx2^-/-^) embryos, respectively.

mirVana microarrays (Ambion) were used to profile microRNA signatures between E16.5 wild-type and homozygous embryos conditional mutants (Mlc2vCre^-^Pitx2^fl/fl^ vs Mlc2vCre^+^Pitx2^-/-^). Tissue samples were collected and pooled from isolated ventricular chambers (n=5) corresponding to E16.5 control (Mlc2vCre^-^Pitx2^fl/fl^) and homozygous conditional (Mlc2vCre^+^Pitx2^-/-^) embryos, respectively 20 μg of total RNA was used to hybridize each microRNA microarray and two distinct microarrays were assessed per developmental stage analyzed. Each microarray contains quadruplicates of mouse/ human/ rat mature microRNAs identified to date (∼500 microRNAs). microRNA ‐Cy3 labeling, microarray hybridization and washing steps were performed according to the manufacturer´s guidelines. Detailed information regarding the hybridization conditions and bioinformatic analyses is given in Supplementary data.

### qRT-PCR

mRNA qRT-PCR was performed in Mx3005Tm QPCR System with an MxPro QPCR Software 3.00 (Stratagene) and SyBR Green detection system. Two internal controls, mouse βactin and GAPDH, were used in parallel for each run. Each PCR reaction was performed at least three times to obtain representative averages. Further detail information is provided in Supplementary data.

microRNA qRT-PCR was performed using Exiqon LNA microRNA qRT-PCR primers and detection kit according to manufacturer’s guidelines. All reactions were always run in triplicates using 5S as normalizing control, as recommended by the manufacturer. Further detail information is provided in Supplementary data.

The Livak method was used to analyze the relative quantification RT-PCR data (28) and normalized in all cases taking as 100% the wild-type (control) value, as previously described (29). The primers used for qRT-PCR amplification of mRNA and microRNAs are listed on **Supplementary Table 1.**

### ECG recordings

Mice were anesthetized with 2mg/Kg Ketamine (Parker-Davis) intraperitoneally. Surface electrocardiograms (ECG) were recorded and analyzed using a digital acquisition and analysis system (Power Lab/4SP; www.adinstrument.com). ECG measurements were performed as previously reported (26-29); additional details are provided in Supplementary Methods.

### Cell culture and Pitx2c transfection assays

HL-1 cells (6 *10^5^ cells per well) were transfected with CMV-Pitx2c construct using lipofectamine 2000 (Invitrogen), according to manufacturer’s guidelines. Similarly, primary cultures of neonatal (2 days post-natal) cardiomyocytes were isolated using standard procedure, cultured accordingly and transfected with CMV-Pitx2c construct using lipofectamine 2000 (Invitrogen), according to manufacturer’s guidelines. Cells were harvested for 48 hours and processed for RNA isolation as previously described. Transfection efficiency was evaluated by assessment of CMV-EGFP transfected cells, which resulted in all cases in more that 60% transfected cells. In addition, in all cases, Pitx2c quantitation was evaluated by qRT-PCR, which resulted in 5 to 8-fold increase.

### Cell culture and microRNA transfection assays

HL-1 cells (6 *10^5^ cells per well) were transfected with Pre-miRs (Ambion, USA) at 50 nM using lipofectamine 2000 (Invitrogen) according to manufacturer’s guidelines. Negative controls included nontransfected cells as well as FAM-labeled pre-miR negative control transfected cells, which also allowed transfection efficiency evaluation. In all cases, transfection efficiencies were greater than 50%, as revealed by observation of FAM-labeled pre-miR transfection. After 4 hours posttransfection, HL-1 cells were cultured in appropriate cell culture media and collected after 48 hours.

### Statistical analysis

qRT-PCR data statistical analyses were performed using unpaired Student t‐ test. Significance level or p values are stated on each corresponding figure legend. One-way ANOVA with Brown-Forsythe post-hoc test was used when appropriate. Group differences were declared significant at p <0.05.

## RESULTS

### Ventricular-specific Pitx2 deletion leads to ventricular hypertrophy and dilation

Pitx2 display heterogenous expression in the developing ventricles (10,23,34,35). We analysed the Pitx2 expression levels in right ventricular free wall (RV), left ventricular free wall (LV) and interventricular septum (IVS) in fetal and adult stages. As depicted in **Supplementary Figure 1**, *Pitx2b* and *Pitx2c* display different expression levels in RV, LV, and IVS at both stages. Importantly, while *Pitx2b* and *Pitx2c* are moderately low in the embryonic RV, enhanced expression is observed in the adult RV. We therefore generated conditional tissue-specific *Pitx2* mutant mice by intercrossing a *Pitx2* floxed mouse line (7) with ventricular-specific Cre (Mlc2vCre) deletor (27) mouse line, rendering ventricular-specific (Mlc2vCre^+^Pitx2^-/-^) *Pitx2* mutant mice, as reported by Chinchilla et al. (23). Deletion of Pitx2 in the ventricular chambers resulted in viable homozygous. qRT-PCR expression analyses of the two Pitx2 isoforms expressed in the adult heart (*Pitx2b* and *Pitx2c*)(35) demonstrated a ∼80% reduction of Pitx2 expression within the ventricular chambers (**Supplementary Figure 2**).

In order to investigate if morphogenetic defects were present in ventricular specific Pitx2 conditional mouse mutants, control (Cre^-^, Pitx2^fl/fl^; i.e Mlc2vCre^-^Pitx2^fl/fl^) and homozygous mutant (Cre^+^Pitx2^-/-^; i.e. Mlc2vCre^+^Pitx2^-/-^) mouse embryos were generated, collected at different developmental stages and morphologically analyzed. Morphological examination of Mlc2vCre^+^Pitx2^-/-^ mutant embryos at E13.5 displays no overt morphological defects (**Figure 1A-B**). At E14.5-E15.5, Mlc2vCre^+^Pitx2^-/-^ embryos display an increased thickness of the interventricular septum (IVS) in 60% of the cases and such morphological defects in the IVS becomes more noticeable with further development (80% in E16.5-E18.5), yet with apparently normal right and left ventricular free walls (**Figure 1C-D**). Importantly, cell proliferation, as revealed by PCNA and phospho-histone 3 labelling experiments, display no significant changes either in the right or left ventricular free walls, or in the IVS (**Figure 1E**) suggesting therefore that the increase in the IVS thickness is due to hypertrophy and not to hyperplasia. In line with these findings, hypertrophic markers such as *Mhy7*, *Mef2c* and *Nppa* are significantly increased in mutant Mlc2vCre^+^Pitx2^-/-^ as compared to age-matched controls (**Figure 1F**), whereas other molecular markers such as *Mlc2a* and *Mlc2v* display no significant differences (data not shown).

**Figure 1.**
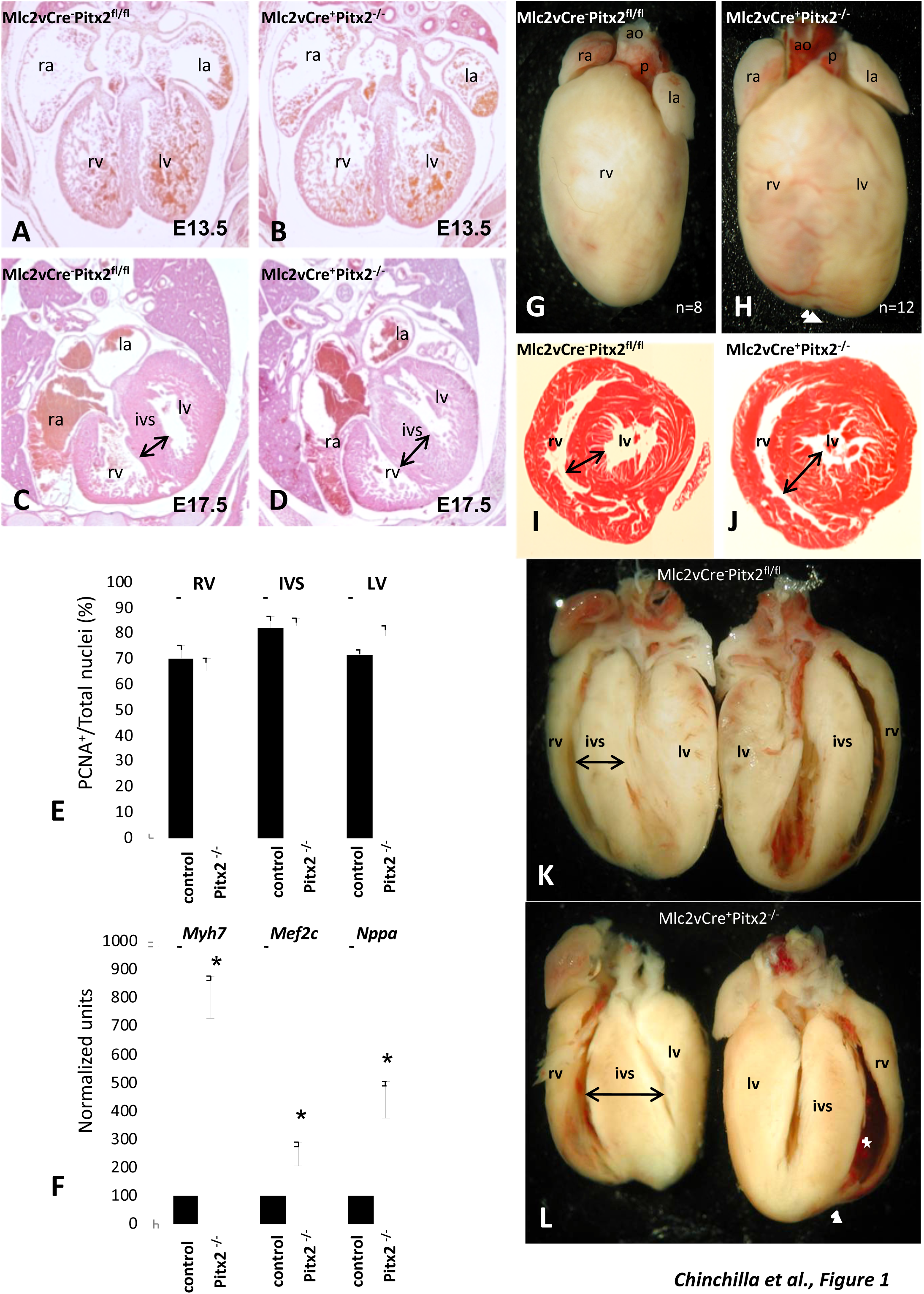
Morphological defects of embryonic and adult ventricular-specific Pitx2 conditional mutants. A-D Histological transverse sections of E13.5 (A,B) and E17.5 (C,D) embryonic hearts corresponding to Mlc2vCre^-^*Pitx2*^flox/flox^ (A,C)(n=6, n=8, respectively) and Mlc2vCre^+^*Pitx2*^-/-^ (B,D)(n=6, n=8, respectively) embryos illustrating similar morphological characteristics at E13.5 (A,B) while an increased thickness of IVS is observed at E17.5 in Mlc2vCre^+^*Pitx2*^-/-^ (C) but not in Mlc2vCre^+^*Pitx2*^flox/flox^ (D) embryos. Panel E illustrate cell proliferation analyses in both left, right myocardium and IVS corresponding to Mlc2vCre^-^*Pitx2*^flox/flox^ (control) and Mlc2vCre^+^*Pitx2*^-/-^ (*Pitx2*^-/-^) E17.5 embryos (n=3). Note that no statistically significant difference in cell proliferation ratio is observed. Panel F illustrates qRT-PCR analyses of several hypertrophy molecular markers at similar developmental stages (E17.5) Observed increased expression of *Myh7*, *Mef2c* and *Nppa* in Mlc2vCre^+^*Pitx2*^-/-^ (*Pitx2*^-/-^) as compared to Mlc2vCre*Pitx2*^flox/flox^ (control) embryos. Panels G-H and K-L whole-mount ventral views (G,H) and longitudinal views (K,L) corresponding to adult Mlc2vCre-*Pitx2*^flox/flox^ (K,G) and Mlc2vCre^+^*Pitx2*^-/-^ (H,L) hearts, respectively. Observe the increased heart size in Mlc2vCre^+^*Pitx2*^-/-^ compared to control Mlc2vCre^-^*Pitx2*^flox/flox^ hearts and partially misshaped (arrowhead, panel H and L). IVS thickness is significantly increased in Mlc2vCre^+^*Pitx2*^-/-^ (L) compared to control Mlc2vCre^-^*Pitx2*^flox/flox^ (G) hearts. In addition, the right ventricular lumen is significantly increased (asterisk, panel L). Transversal histological sections of adult ventricular Mlc2vCre-*Pitx2*^flox/flox^ (I) and Mlc2vCre^+^*Pitx2*^-/-^ (J) chambers illustrate a significant IVS thickness (double arrows). Experiments on panel F were performed in three independent biological samples of pooled embryonic hearts (≥3 hearts per condition) (n≥9). Represented values correspond to the average mean ± sd. *p <0.01

Beside the increased in IVS thickness, Mlc2vCre^+^Pitx2^-/-^ mutants embryos did not display any other overt morphological defects during embryogenesis and Mlc2vCre^+^Pitx2^-/-^ mice are viable until adulthood. Mlc2vCre^+^Pitx2^-/-^ adult hearts display dysmorphogenic ventricular chambers with an engrossed IVS (∼83%; 10/12), as illustrated in **Figure 1G-L**, and dilation of the right ventricular chamber (∼41%; 5/12) as depicted in **Figure 1K-L.** Curiously, no overt heart-to-body weight ratio significant differences were observed (controls, n=30, 7,18±1,24; Mlc2vCre^+^Pitx2^-/-^, n=20, 6,80±1,31).

### Pitx2 loss-of-function leads QRS prolongation

We have recently demonstrated that atrial chamber-specific Pitx2 insufficiency leads to cellular and electrophysiological changes prone to develop atrial arrhythmias (23), supporting GWAS studies that linked PITX2 and atrial fibrillation (11-20). We determine herein if ventricular-specific Pitx2 insufficiency also leads to electrophysiological impairment. ECG recordings on adult Mlc2vCre^+^Pitx2^-/-^ hearts demonstrate no significant differences in heart rate (HR) or RR interval as compared to controls (**Figure 2A-C**). Interestingly, prolongation of the QRS, QT and PR intervals (**Figure 2C**) was identified in Mlc2vCre^+^Pitx2^-/-^ adult hearts (**Figure 2B, B´, B´´**) as compared to controls (**Figure 2A, A´, A´´**). Widening of the QT interval is mainly due by either impaired depolarization/repolarization of ventricular excitation-contraction, and/or by concomitant ventricular conduction system defects, which can be further amplified by the presence of cardiac hypertrophy (36-40). Morphological examination of the ventricular conduction system displays similar distribution of the distinct components of the ventricular conduction system in Mlc2vCre^+^Pitx2^-/-^ adult hearts (**Figure 2G-I**) as compared to wild-type age-matched controls (**Figure 2D-F**)(**Supplementary Figure 2**). In some hearts (40%; 2/5), nonetheless insulation of the His bundle is unequally developed (**Figure 2G-H**), which might related to the prolonged PR interval observed in these mice, yet other unknown causes might also account for such electrophysiological alteration.

**Figure 2.**
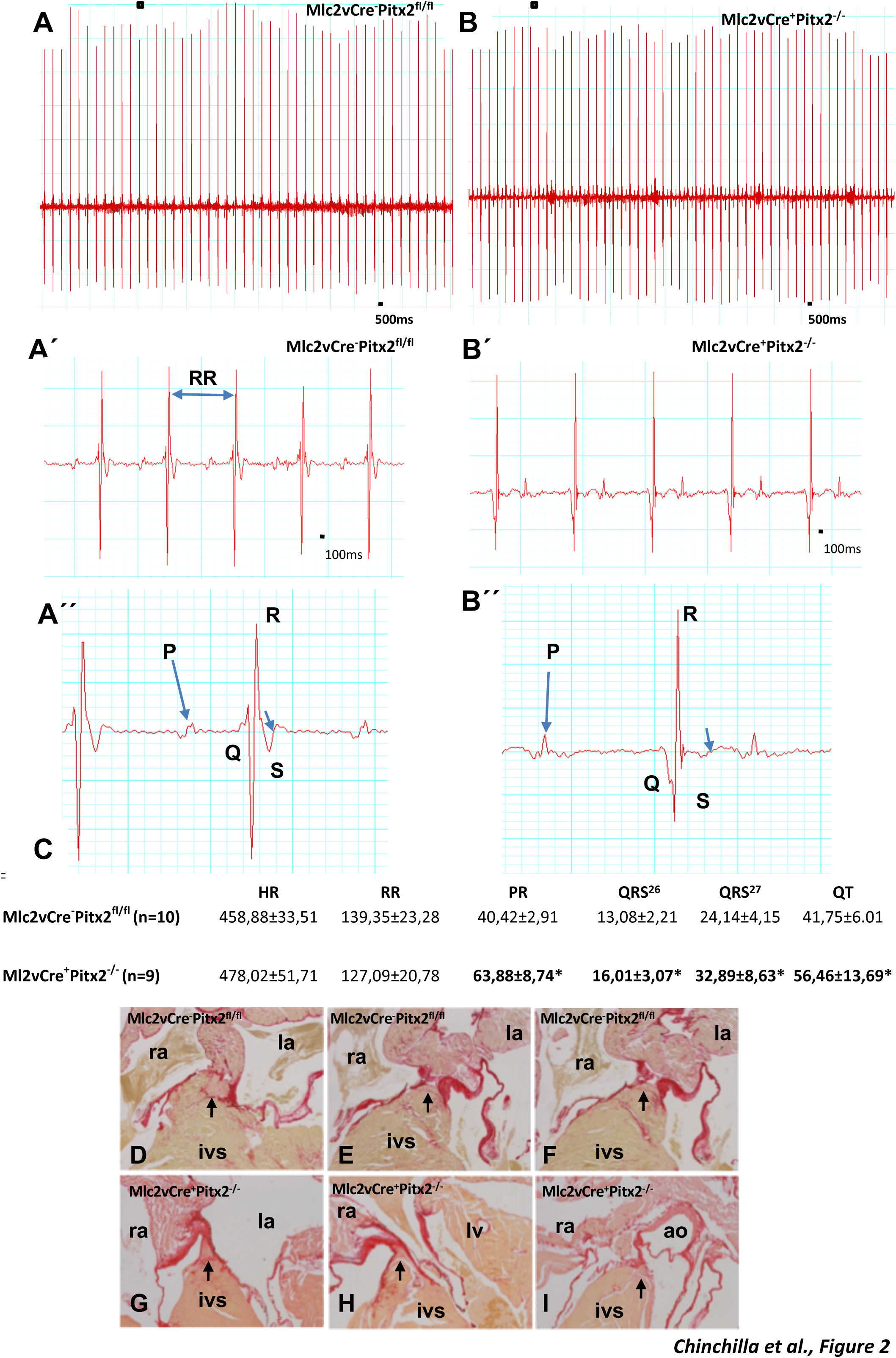
ECG measurements and CCS morphology in adult ventricular-specific Pitx2 conditional mutants. Representative electrocardiogram (ECG) recordings (A-B) of the adult hearts corresponding to Mlc2v^-^Cre*Pitx2*^flox/flox^ (A, A´, A´´) and Mlc2vCre^+^*Pitx2*^-/-^ (B, B´, B´´) mice. A´´ and B´´ are close-ups of A´ and B´, respectively. Panel C displays the statistical significance of HR, RR, PR, QRS and QT intervals between Mlc2vCre^-^*Pitx2*^flox/flox^ and Mlc2vCre^-^*Pitx2*^-/-^ adult mice. Ventricular chamber-specific Pitx2 conditional mutants display prolongation of the PR, QRS and QT intervals, while HR and RR intervals are not significantly different. Panels D-I Red Sirius staining of histological sections corresponding to Mlc2vCre^-^*Pitx2*^flox/flox^ (D-F)(n=8) and Mlc2vCre^+^*Pitx2*^-/-^ (G-I)(n=12) adult hearts demonstrating the fibrous tissue deposition within the ventricular conduction system. Note that morphological examination of the conduction system reveals no significant differences with the morphological distribution. Only in some cases (∼40%) insulation of the His bundle is significantly reduced in Mlc2vCre*Pitx2*^-/-^ adult mouse hearts compared to controls.*p <0.01

We have further explored if such electrophysiological defects can be caused by altered expression of those genes encoding ion channels that are genetically associated to long QT syndrome in humans (38) and/or those contributing to similar electrophysiological characteristics in mice (39), given the overt species-specific differences (37-40). As depicted in **Figure 3A**, no differences in expression are observed for *Scn5a* and *Scn1b* in both controls and Pitx2 conditional mutants. However, potassium ion channels involved in the repolarization phase (*Kcnq1* and *Kcnh2*) and resting membrane potential (*Kcnj2*) of the cardiac action potential are significantly decreased in Mlc2vCre^+^Pitx2^-/-^ mouse adult hearts as compared to controls. No significant changes were observed for *Kcne1*, *Kcne2* (data not shown), *Kncj12* and *Kcnd2* (**Figure 3A**) and only a mild decreased was documented for *Kcnd3* (**Figure 3A**). Surprisingly, *Kcna5* was significantly increased (**Figure 3A**). Overall, these data suggest that impaired repolarization underlies the prolonged QT interval reported in this mutant model, while depolarization is unaltered, although we cannot exclude that part of the observed ion channel remodeling could be secondary to the hypertrophic features observed in Mlc2vCre^+^Pitx2^-/-^ mouse adult hearts.

**Figure 3.**
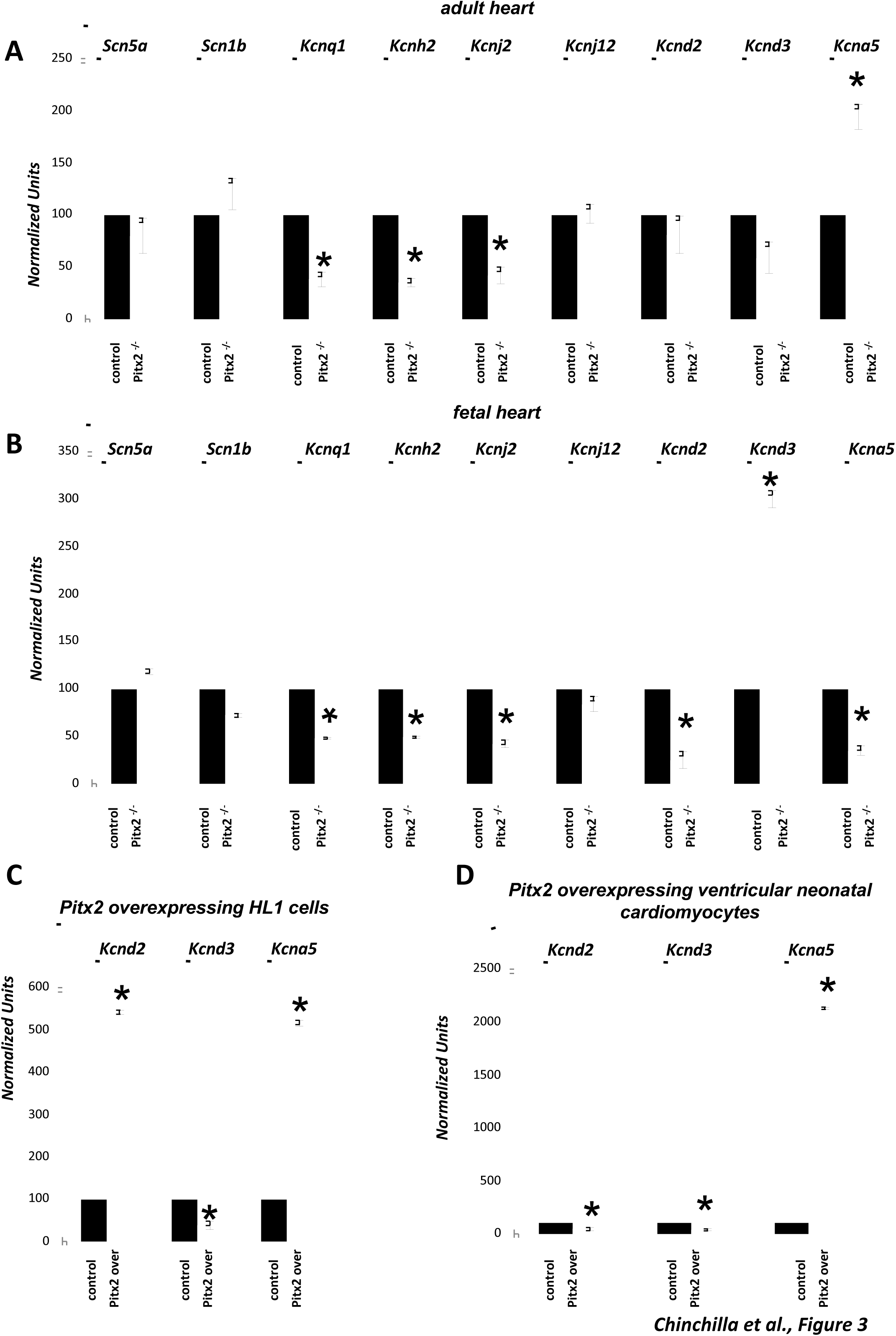
Molecular remodeling of adult ventricular-specific Pitx2 conditional mutants. Panel A illustrates qRT-PCR expression analysis of *Scn5a, Scn1b, Kcnq1, Kcnh2, Kcnj2, Kcnj12, Kcnd2, Kcnd3* and *Kcna5* in ventricular chambers corresponding to adult ventricular-specific Pitx2 conditional hearts (white bars) as compared to controls (black bars). Observe that *Kcnh2, Kcnq1*, *Kcnj2* display significantly decreased expression levels in ventricular chambers while *Kcna5* display increased expression levels. No changes are observed for *Scn1b*, *Scn5a, Kcnj12* and *Kcnd2*. Moderate decreased levels are observed for *Kcnd3,* yet without reaching significant differences. Panel B illustrates qRT-PCR analyses of ion channel expression at fetal stages (E16.5) between Mlc2vCre-*Pitx2*^flox/flox^ (control) and Mlc2vCre^+^*Pitx2*^-/-^ (*Pitx2*^-/-^) ventricular chambers. Note that *Kcnh2, Kcnq1,Kcnj2, Kcnd2* and *Kcna5* expression is significantly decreased in E16.5 Pitx2 conditional hearts (white bars) as compared to controls (black bars) while *Kcnd3* is significantly increased. Panel C illustrates qRT-PCR analyses of *Kcnd2*, *Kcnd3* and *Kcna5* expression in HL-1 atrial cardiomyocytes transfected with Pitx2 (white bars) as compared to non-transfected controls (black bars). Note that *Kcnd2* and *Kcna5* increase their expression after Pitx2 transfection while *Kcnd3* decreases. Panel D illustrates qRT-PCR analyses of *Kcnd2*, *Kcnd3* and *Kcna5* expression in primary culture of neonatal cardiomyocytes transfected with Pitx2 (white bars) as compared to non-transfected controls (black bars). Note that *Kcnd2* and *Kcnd3* decrease their expression after Pitx2 transfection while *Kcna5* increases. All experiments were performed at least three times within three independent biological samples (n=9). Represented values correspond to the average mean ± sd.*p <0.01

To get insights if ion channel remodeling in the adulthood was occurring in Mlc2vCre^+^Pitx2^-/-^ mice, we analysed by qPCR these genes at fetal stages. *Scn5a* and *Scn1b* are similarly expressed in Mlc2vCre^+^Pitx2^-/-^ mouse E16.5 ventricular chambers as compared to age-matched controls. Importantly, potassium channels such as *Kcnq1*, *Kcnh2* and *Kcnj2*, but not *Kcnj12*, are significantly decreased in Mlc2vCre^+^Pitx2^-/-^ mouse E16.5 ventricular chambers as compared to age-matched controls (**Figure 3B**). In addition, *Kcnd2* and *Kcna5* are significantly decreased while *Kcnd3* is significantly increased in Mlc2vCre^+^Pitx2^-/-^ mouse E16.5 ventricular chambers as compared to age-matched controls (**Figure 3B**). Thus, electrophysiological defects observed in the Mlc2vCre^+^Pitx2^-/-^ adult hearts are likely to be primary caused directly by impaired Pitx2 expression in the ventricular chambers. To further sustain the regulatory role of Pitx2 modulating the potassium channel expression we over-expressed Pitx2 in HL-1 cardiomyocytes as well as in primary cultures of ventricular fetal cardiomyocytes. Pitx2 overexpression leads to increased expression of *Kcnd3* and a significantly decreased of *Kcna5* in both HL-1 and primary cultures of fetal cardiomyocytes (**Figure 3C-D**) in line with our findings in Mlc2vCre^+^Pitx2^-/-^ mutant hearts. Surprisingly, Pitx2 over-expression increases *Kcnd2* expression in HL-1 cells, but not in neonatal cardiomyocytes, probably reflecting developmental specie-specific balanced expression of *Kcnd2* and *Kcnd3* in different myocardial settings (38-40). Nonetheless, these data suggest a pivotal role of Pitx2 regulating potassium channel expression in the developing and adult heart, and supports the notion that Pitx2-mediated electrophysiolgical defects precede the hypertrophic features observed in Mlc2vCre^+^Pitx2^-/-^ mutant mice, in line with findings reported in a experimental mouse model of complete AV block (40).

### Pitx2 mediated signalling pathways

The morphogenetic role of Pitx2 has been extensively studied (5-10) and more recently the molecular targets of this transcription factor are progressively emerging (21-25). We therefore decided to get further insights into the molecular pathways controlled by Pitx2 in the developing ventricular chambers, by analyzing the mRNA and microRNA expression profiles by microarrays in E16.5 Mlc2vCre^+^Pitx2^-/-^ embryos. Considering a fold-change larger than ±1.5 and p-value lower than 0.05, 117 mRNAs were up-regulated and 64 were down-regulated **(Supplementary Table 2)**; a subset of the most relevant genes for cardiac development and function is listed on **Figure 4A-B**. Validation of a subset of these mRNAs is provided in **Figure 4C-D**. To further sustain the role of Pitx2 on the regulation of these mRNAs, we over-expressed Pitx2c in HL-1 atrial fetal cardiomyocytes and quantified the expression levels of these mRNAs (**Figure 4E-F**). As observed in **Figure 4E and 4F**, overexpression of Pitx2c is sufficient to induce expression of *Myh11*, *Pcolce2*, *Matn2*, *Nppc* and *Snai2* and to repress *Clcnk*, further validating our mRNA microarray data.

**Figure 4.**
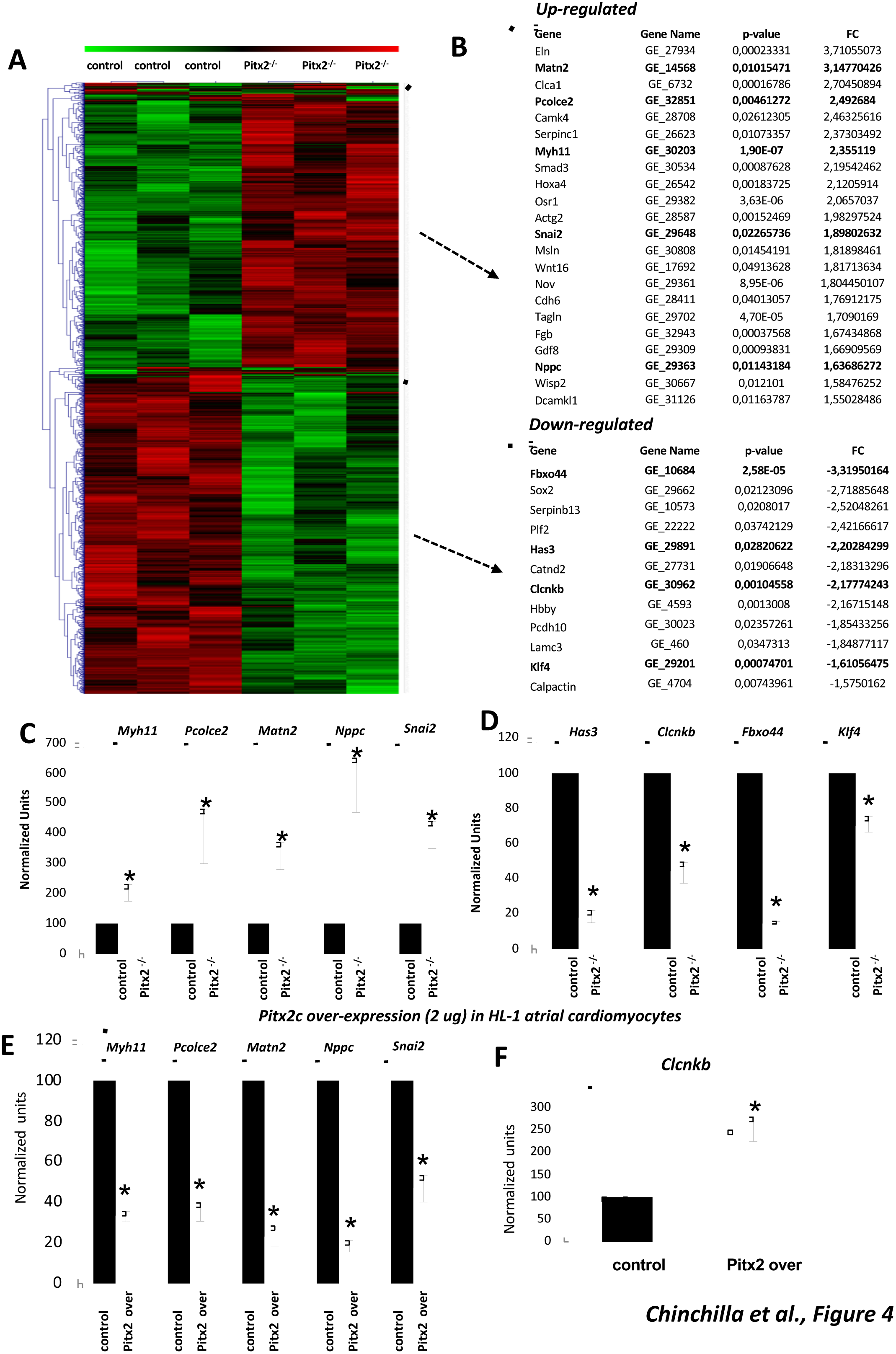
Hierarchical clustering of mRNA microarray expression profiles of E16.5 ventricular-specific Pitx2 conditional mutants. Panel A shows a hierarchical clustering of mRNA microarray expression profiles between E16.5 ventricular-specific Pitx2 conditional mutants compared to controls; only those displaying significant differences (p-values <0.05) are illustrated (green, high; red, low). Panel B shows representative mRNAs with previously reported cardiac expression, selected from those mRNAs differentially expressed; full list of mRNAs is supplied in ***Supplementary Table 2***. Panels C and D illustrate representative mRNAs with increased or decreased expression levels in Mlc2vCre^+^*Pitx2*^-/-^ as compared to Mlc2vCre^-^*Pitx2*^flox/flox^ hearts, which have been validated by qRT PCR, respectively. Panel E and F display qRT-PCR analysis of *Myh11, Pcolce2, Matn2, Nppc, Snai2* (panel E) and *Clcnk* (panel F) expression using Pitx2c over-expressing HL-1 atrial cardiomyocytes. Note that *Pitx2c* over-expression decreases transcript expression of *Myh11, Pcolce2, Matn2, Nppc, Snai2* (panel A) and increases transcript expression of *Clcnk* (panel B), suggesting a regulatory role for Pitx2c modulating the expression of these transcripts. All experiments were performed at least three times within three independent biological samples (n=9). Represented values correspond to the average mean ± sd.*p <0.01

Similarly, we conducted a microRNA microarray expression profiling at E16.5 comparing Mlc2vCre^+^Pitx2^-/-^ hearts with wild type control. We identified nine up-regulated (*miR-1, miR-7, miR-26b, miR-148a, miR-199, miR-299, miR-328, miR-350, let-7f*) and six down-regulated (*miR-202, miR-344, miR-363, miR-379, miR-409, miR-499*) microRNAs. qRT-PCR analyses validated our microarray analyses as depicted in **Figure 5A-B**. In addition, Pitx2c overexpression in HL-1 cardiomyocytes is sufficient to induce miR-202, but not miR-344 or miR-409 (**Figure 5C**) and to decrease the expression levels of miR-1, miR-148, miR-199, miR-328 but not miR-7 (**Figure 5D**). Thus, these data demonstrate that Pitx2 plays a pivotal role regulating the expression of a subset of microRNAs in the developing myocardium.

**Figure 5.**
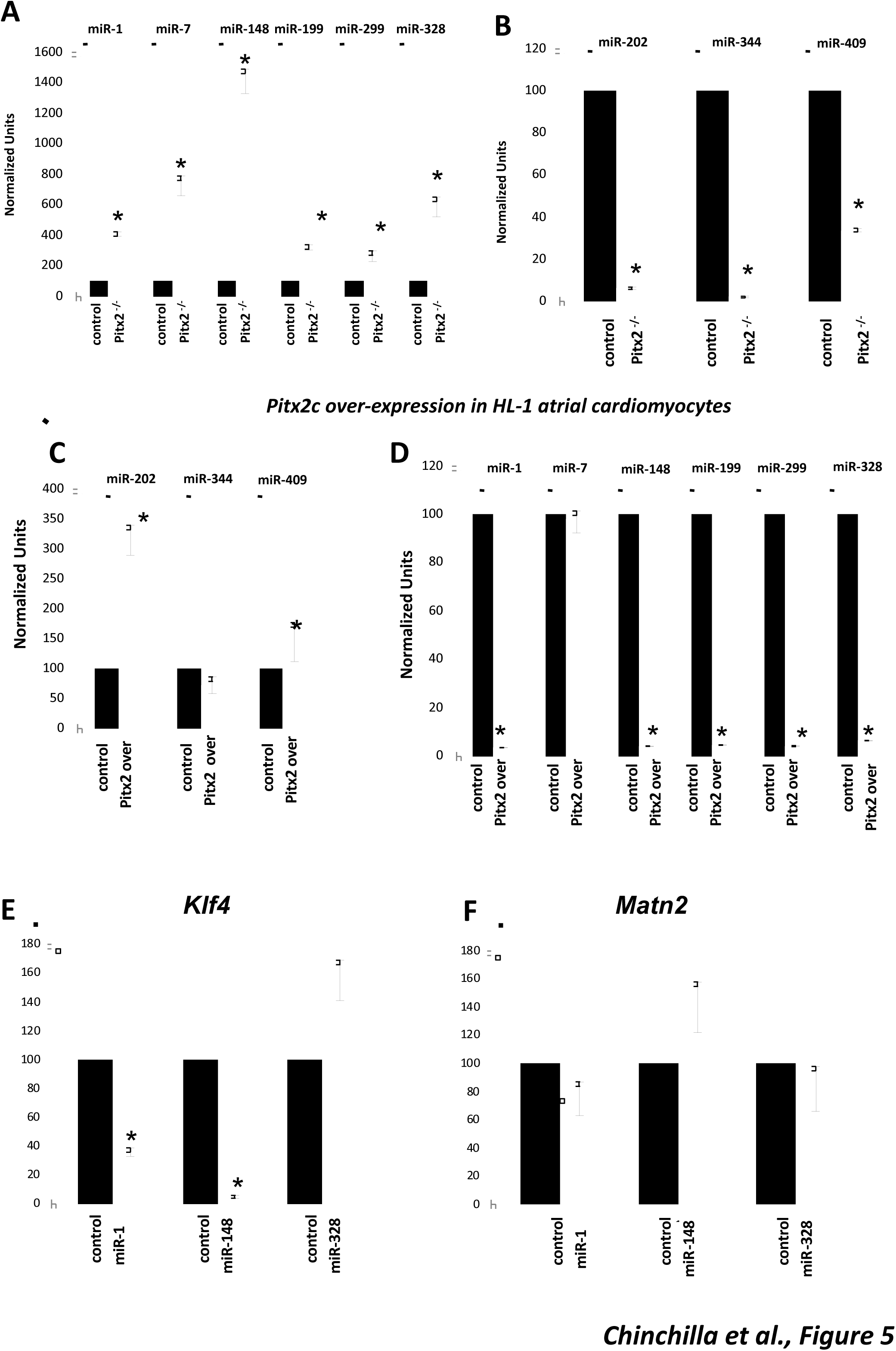
microRNA expression profiling of E16.5 ventricular-specific Pitx2 conditional mutants. Panels A and B illustrates qRT-PCR microarray validated microRNAs with increased (*miR-1, miR-7, miR-148, miR-199, miR-299, miR-328;* panel A) or decreased (*miR-202, miR-344, miR-409*; panel B) expression in Mlc2vCre^+^*Pitx2*^-/-^ as compared to Mlc2vCre^-^*Pitx2*^flox/flox^ embryos. qRT-PCR analysis of *miR-1*, *miR-7*, *miR-148*, *miR-199*, *miR-299* and *miR-328* (panel C) and *miR-202*, *miR-344* and *miR-409* (panel D) expression in Pitx2c over-expressing HL-1 in atrial cardiomyocytes. Note that Pitx2c over-expression decreases transcript expression of *miR-202*, *miR-344* and *miR-409* (panel C) and increases transcript expression of *miR-1*, *miR-7*, *miR-148*, *miR-199*, *miR-299* and *miR-328* (panel B), supporting that Pitx2c displays a regulatory role on the expression of these genes. Panel E and F illustrate qRT-PCR analyses of Klf4 (panel E) and Matn2 (panel F) expression levels in HL-1 cardiomyocytes transfected with miR-1, miR-148 and miR-328, respectively, as compared to nontransfected controls. All experiments were performed at least three times within three independent biological samples (n=9). Represented values correspond to the average mean ± sd.*p <0.01.

Interestingly several mRNAs which display impaired expression in ventricular-specific Pitx2 insufficient mice are reported to be putative targets of discrete microRNAs deregulated in Mlc2vCre^+^Pitx2^-/-^ embryos. In particular, we noticed that the cardiac hypertrophy-related transcription factor (Klf4) (41) is targeted by miR-1 (42) and putatively by miR-148 but not by miR-328 as revealed by TargetScan algorithm. We therefore explore if overexpression of these microRNAs would result in impaired expression of Klf4. HL1 cardiomyocytes transfection assays revealed that miR-1 and miR-148, but not miR-328, leads to severe *Klf4* downregulation (**Figure 5E**), without altering for *Matn2* (**Figure 5F**), another Pitx2 deregulated mRNA, demonstrating a key role of these microRNAs in the regulation of Klf4.

Given the fact that Pitx2 display heterogeneous expression in the developing ventricles as reported in this study, we subsequently tested whether *miR-1* or *miR-328* display similar or complementary Pitx2 expression patterns in distinct adult ventricular regions. Interestingly, *miR-1* and *miR-328* display complementary patterns, further supporting the regulatory role of Pitx2 on these genes (**Supplementary Figure 3**).

## DISCUSSION

### Pitx2 insufficiency in the developing ventricular myocardium leads to IVS thickening

*Pitx2* is a homeobox transcription factor with a relevant role during cardiogenesis (2-9). Seminal papers have demonstrated that lack of *Pitx2* in the developing embryo compromised cardiac morphogenesis (7, 8). More recently, it has been demonstrated that such roles are partially executed in fully mature cardiomyocytes (10). We have generated chamber-specific conditional *Pitx2* mutants with reduced 50-60% expression of Pitx2b and Pitx2c, respectively, as consequence of incomplete Cre recombination in the developing atrial and ventricular chambers (23, 25, 43), since both isoforms are expressed at fetal and adult stages (35). In this setting, *Pitx2* insufficiency in the atrial chambers (23) and/or the ventricular chambers (this study) displays mild embryonic morphogenetic defects that are compatible with life. Atrial *Pitx2* chamber-specific mutants display atrial dilatation (23) while ventricular *Pitx2* chamber-specific mutants displayed an increase on the IVS thickness. These data demonstrate therefore a chamber-specific roIe for Pitx2 in cell cycle regulation since in atrial *Pitx2* insufficiency leads to up-regulation of *Bmp10* with concomitant increase in cell rate proliferation as well as in several cell cycle regulator genes, such as *cyclin D1*, *cyclin D2* and *cmy*c (23), whereas in the ventricular chambers, cell cycle is not increased, and therefore the thickening of the IVS occurs at the expense of increasing the cell size, i.e. hypertrophy, a notion further supported by increased expression of hypertrophic markers such as *Nppa*, *Mef2c* and *Mhy7*. These findings shed the lights on previous controversial findings arguing that Pitx2 distinctly increases (44-47), decreases (48-49) or has no effect (21) cell proliferation, suggesting therefore a tissue-dependent of Pitx2 on cell cycle regulation.

### Pitx2 insufficiency alters ion channel function in the adult ventricular myocardium leading to an increased QT interval

Recent GWAS in humans have provided evidence of several risk variants located near to PITX2 are highly associated with AF (11-21), supporting a role of PITX2 in AF. Pitx2 atrial-specific insufficiency mice display cellular, molecular and electrophysiological defects that are prone to provoke atrial arrhythmias (23,25), in line with other reports (21, 22, 24). Interestingly, ventricular chamber-specific Pitx2 conditional mutants also present electrophysiological defects as revealed by ECG measurements; both QRS and QT intervals are prolonged. The amplitude of the QT segment is determined by the interval between the depolarization and repolarization of the ventricular chambers (33), and thus modulated by the conduction propagation and/or the inherent depolarization/repolarization characteristics of the ventricular myocytes. QRS duration, independently as whether defined as the end of the J wave (31) or by the intersection of the S wave with the isoelectric line (30), underestimates the ventricular conduction assessment in mice (33). Importantly, morphological examination of the ventricular conduction system reveals no overt alterations, suggesting thus that QT widening is mainly caused by impaired ventricular depolarization/repolarization. In addition, it is important to highlight that Pitx2 ventricular conditional mouse adult hearts displayed increased PR interval, suggesting that a conduction delay in the AVN might be also contributing to the electrophysiological defects recorded in these mice. Interestingly, similar electrophysiological defects have been recently reported by Tao et al. (24) in mice with conditional deletion of Pitx2 in the adult myocardium, yet the underlying molecular mechanisms, in both models, remains to be fully elucidated

We next analysed the molecular determinants of cardiac depolarization and repolarization. No significant differences were observed for the major determinants of depolarization (*Scn5a, Scn1b*).Importantly, significant decrease on *Kcnh2* and *Kcnq1*, which might decrease I_Kr_ and I_Ks_ currents were observed in both fetal and adult hearts. These data are in line with recent reports demonstrating a Pitx2 regulation of I_Kr_ and I_Ks_ in HL1 cardiomyocytes (50). Point mutations in KCNH2 and KCNQ1 have been reported in LQT1 and LQT2 syndromes, respectively (51). Interestingly, in mice and rabbits, functional suppression of these channels have been previously reported to cause long QT (52, 53). Thus, these data suggest that our model can be regarded as a long QT-like mouse mutant model, in line with findings in humans (51, 54). However, it is important to mention that repolarization phase in mice is also dependent of additional currents, such as I_TO_ (*Kcnd2*, *Kcnd3*) and I_Kur_ (*Kcn5a*). In this context, it is important to highlight that *Kcnd2*, *Kcnd3* and *Kcn5a* are impaired in Pitx2 ventricular specific mutant at fetal stages. Curiously, only increased Kcn5a is observed at adult stages, demonstrating partially divergent expression pattern in fetal and adult stages. These findings might be regarded as secondary ion channel remodeling contributing to the adult Mlc2vCre^+^Pitx2^-/-^ phenotype. In addition, impaired expression of determinants of the resting membrane potential (*Kcnj2*) are also observed in both fetal and adult stages. Thus these data underscore the notion that Pitx2 modulates multiple pathways contributing to the repolarization phase in the ventricular chambers that, if impair could lead to long QT. In this setting, Pitx2 over-expression experiments in HL1 cells and primary neonatal ventricular cardiomyocytes futher underscore this notion. Overall, our data demonstrate that Pitx2 plays a crucial role controlling several electrophysiological parameters not only in the atrial (22), but also in the ventricular myocardium, which are prone to evoke arrhythmias. Secondly, our data suggest that Pitx2 impairment leads primarily to ion channel remodeling, yet it remains to be elucidated whether cardiac hypertrophy is cause or consequence of these abnormalities.

### Pitx2 regulates miRNA-mRNA interactions leading to cardiac hypertrophy and arrhythmias

Genome-wide gene expression profiling by means of microarray analyses, and more recently by ChIP-seq (24), have provided insights into the molecular mechanisms underlying the cellular and molecular defects observed in the Pitx2 mouse mutants (22-25). Kirchhof et al. (22) recently described mRNA differential expression profile in left and right atrial chambers of Pitx2c deficient mice. These authors reported a significant differential expression of calcium ion binding genes, gap and tight junctions and calcium and potassium channels. Our results using a similar approach in our Pitx2 ventricular-specific insufficient mice indicate that Pitx2 preferentially regulates signaling pathways such as mesoderm development, cell-cell signaling, stemness and cell growth (***Supplementary Table 4***), suggesting therefore that distinct regulatory networks are controlled by Pitx2 in the atrial *vs* ventricular chamber.

In addition to genome-wide mRNA gene expression analyses we have also performed microRNA comparative analyses in fetal Pitx2 ventricular-specific insufficient mice. We have identified a subset of microRNAs regulated by Pitx2, as underscored by microarray, qRT-PCR validation and Pitx2c overexpression experiments in HL-1 atrial cardiomyocytes. Importantly a distinct set of microRNAs are impaired in Pitx2 atrial-specific mouse mutants, contributing to ion channel dysregulation in the atrial chambers (25,55). In the ventricular context, it is important to highlight three microRNAs; miR-1, miR-148 and miR-328. Pitx2 is necessary and sufficient to regulate miR-1, miR-148 and miR-328 expression. miR-1 null mutant mice display to conduction disturbances (56) and miR-1 over-expression regulates *Kcnj2* expression (57, 58), providing therefore a direct regulatory link between for Pitx2 >miR-1 >Kcnj2 in arrhythmogenesis. Moreover, impaired expression of miR-328 in our Pitx2 ventricular-specific insufficient mice further underscores the molecular link between Pitx2 and arrhythmias, since forced expression of miR-328 in the heart leads to atrial fibrillation (59). In addition, miR-1 has been reported to regulate Kcnj2 (57, 58) as well as Klf4 (41), a transcription factor that critically regulates cardiac hypertrophy (40). We provide herein evidences that not only miR-1 (41), but also miR148 can also modulate Klf4 expression, supporting therefore that the hypertrophic phenotype observed in our ventricular-specific Pitx2 mouse mutants can be driven by Pitx2 impaired expression of miR-1 and miR-148, impacting on Kfl4 expression and thus resulting in hypertrophy.In summary our data demonstrate that Pitx2 insufficiency in the ventricular chamber leads to cellular and molecular remodeling, mediated in part by microRNAs, which results in hypertrophic ventricular chambers and electrophysiological defects resembling long QT.

## ACKNOWLEDGMENTS

We would like to thank Phil Gage (University of Michigan Medical School, Ann Harbor, Michigan, USA) and Kenneth Chien (University of California, San Diego, USA) for reagents. This work is partially supported by the VI EU Integrated Project “Heart Failure and Cardiac Repair” LSHM-CT-2005-018630 to DF, a grant from the Junta de Andalucía Regional Council to DF (CVI-6556), a grant from the Junta de Andalucía Regional Council to AA (CTS-03878) and grants from the Ministry of Science and Innovation of the Spanish Government to DF (MICINN BFU2009-11566) and to AA (MICINN BFU-2008-01217). This work is partially supported by a translational CNIC grant (2009-08) to DF. This work is partially supported by a grant from the University of Jaén (UJA2009/12/11) to JND.

## REFERENCES

1. Burdine RD, Schier AF. Conserved and divergent mechanisms in left-right axis formation. Genes Dev. 2000;14:763–776.

2. Schweickert A, Campione M, Steinbeisser H, Blum M. Pitx2 isoforms: involvement of Pitx2c but not Pitx2a or Pitx2b in vertebrate left-right asymmetry. Mech Dev. 2000;90:41–51.

3. Campione M, Steinbeisser H, Schweickert A, Deissler K, van Bebber F, Lowe LA, Nowotschin S, Viebahn C, Haffter P, Kuehn MR, Blum M. The homeobox gene Pitx2: mediator of asymmetric left-right signaling in vertebrate heart and gut looping. Development. 1999;126:1225–1234.

4. Campione M, Ros MA, Icardo JM, Piedra E, Christoffels VM, Schweickert A, Blum M, Franco D, Moorman AF. Pitx2 expression defines a left cardiac lineage of cells: evidence for atrial and ventricular molecular isomerism in the iv/iv mice. Dev Biol. 2001;231:252–264.

5. Logan M, Pagán-Westphal SM, Smith DM, Paganessi L, Tabin CJ. The transcription factor Pitx2 mediates situs-specific morphogenesis in response to left-right asymmetric signals. Cell. 1998; 94:307–317.

6. Piedra ME, Icardo JM, Albajar M, Rodriguez-Rey JC, Ros MA. Pitx2 participates in the late phase of the pathway controlling left-right asymmetry. Cell. 1998;94:319–324.

7. Gage PJ, Suh H, Camper SA. Dosage requirement of Pitx2 for development of multiple organs. Development. 1999;126:4643–4651.

8. Ai D, Liu W, Ma L, Dong F, Lu MF, Wang D, Verzi MP, Cai C, Gage PJ, Evans S, Black BL, Brown NA, Martin JF. Pitx2 regulates cardiac left-right asymmetry by patterning second cardiac lineage-derived myocardium. Dev Biol. 2006;296:437–449.

9. Ammirabile G, Tessari A, Pignataro V, Szumska D, Sutera Sardo F, Benes J Jr, Balistreri M, Bhattacharya S, Sedmera D, Campione M. Pitx2 confers left morphological, molecular, and functional identity to the sinus venosus myocardium. Cardiovasc Res.. 2012; 93(2):291–301.

10. Tessari A, Pietrobon M, Notte A, Cifelli G, Gage PJ, Schneider MD, Lembo G, Campione M. Myocardial Pitx2 differentially regulates the left atrial identity and ventricular asymmetric remodeling programs. Circ Res. 2008;102:813–822.

11. Gudbjartsson DF, Arnar DO, Helgadottir A, Gretarsdottir S, Holm H, Sigurdsson A, Jonasdottir A, Baker A, Thorleifsson G, Kristjansson K, Palsson A, Blondal T, Sulem P, Backman VM, Hardarson GA, Palsdottir E, Helgason A, Sigurjonsdottir R, Sverrisson JT, Kostulas K, Ng MC, Baum L, So WY, Wong KS, Chan JC, Furie KL, Greenberg SM, Sale M, Kelly P, MacRae CA, Smith EE, Rosand J, Hillert J, Ma RC, Ellinor PT, Thorgeirsson G, Gulcher JR, Kong A, Thorsteinsdottir U, Stefansson K. Variants conferring risk of atrial fibrillation on chromosome 4q25. Nature 2007;448:353–357.

12. Kääb S, Darbar D, van Noord C, Dupuis J, Pfeufer A, Newton-Cheh C, Schnabel R, Makino S, Sinner MF, Kannankeril PJ, Beckmann BM, Choudry S, Donahue BS, Heeringa J, Perz S, Lunetta KL, Larson MG, Levy D, MacRae CA, Ruskin JN, Wacker A, Schömig A, Wichmann HE, Steinbeck G, Meitinger T, Uitterlinden AG, Witteman JC, Roden DM, Benjamin EJ, Ellinor PT. Large scale replication and meta-analysis of variants on chromosome 4q25 associated with atrial fibrillation. Eur Heart J. 2009;30:813–819.

13. Franco D, Christoffels VM, Campione M. Homeobox transcription factor Pitx2: The rise of an asymmetry gene in cardiogenesis and arrhythmogenesis. Trends Cardiovasc Med. 2014 Jan;24(1):23–31

14. Delaney JT, Jeff JM, Brown NJ, Pretorius M, Okafor HE, Darbar D, Roden DM, Crawford DC. Characterization of genome-wide association-identified variants for atrial fibrillation in African Americans. PLoS One. 2012;7(2):e32338.

15. Henningsen KM, Olesen MS, Haunsoe S, Svendsen JH. Association of rs2200733 at 4q25 with early onset of lone atrial fibrillation in young patients. Scand Cardiovasc J. 2011 Dec;45(6):324–6.

16. Virani SS, Brautbar A, Lee VV, Elayda M, Sami S, Nambi V, Frazier L, Wilson JM, Willerson JT, Boerwinkle E, Ballantyne CM. Usefulness of single nucleotide polymorphism in chromosome 4q25 to predict in-hospital and long-term development of atrial fibrillation and survival in patients undergoing coronary artery bypass grafting. Am J Cardiol. 2011 May 15;107:1504–9.

17. Lee KT, Yeh HY, Tung CP, Chu CS, Cheng KH, Tsai WC, Lu YH, Chang JG, Sheu SH, Lai WT. Association of RS2200733 but not RS10033464 on 4q25 with atrial fibrillation based on the recessive model in a Taiwanese population. Cardiology. 2010;116(3):151–6.

18. Husser D, Adams V, Piorkowski C, Hindricks G, Bollmann A. Chromosome 4q25 variants and atrial fibrillation recurrence after catheter ablation. J Am Coll Cardiol. 2010, 23;55(8):747–53.

19. Body SC, Collard CD, Shernan SK, Fox AA, Liu KY, Ritchie MD, Perry TE, Muehlschlegel JD, Aranki S, Donahue BS, Pretorius M, Estrada JC, Ellinor PT, Newton-Cheh C, Seidman CE, Seidman JG, Herman DS, Lichtner P, Meitinger T, Pfeufer A, Kääb S, Brown NJ, Roden DM, Darbar D. Variation in the 4q25 chromosomal locus predicts atrial fibrillation after coronary artery bypass graft surgery. Circ Cardiovasc Genet. 2009 Oct;2(5):499–506.

20. Lubitz SA, Lunetta KL, Lin H, Arking DE, Trompet S, Li G, Krijthe BP, Chasman DI, Barnard J, Kleber ME, Dörr M, Ozaki K, Smith AV, Müller-Nurasyid M, Walter S, Agarwal SK, Bis JC, Brody JA, Chen LY, Everett BM, Ford I, Franco OH, Harris TB, Hofman A, Kääb S, Mahida S, Kathiresan S, Kubo M, Launer LJ, Macfarlane PW, Magnani JW, McKnight B, McManus DD, Peters A, Psaty BM, Rose LM, Rotter JI, Silbernagel G, Smith JD, Sotoodehnia N, Stott DJ, Taylor KD, Tomaschitz A, Tsunoda T, Uitterlinden AG, Van Wagoner DR, Völker U, Völzke H, Murabito JM, Sinner MF, Gudnason V, Felix SB, März W, Chung M, Albert CM, Stricker BH, Tanaka T, Heckbert SR, Jukema JW, Alonso A, Benjamin EJ, Ellinor PT. Novel genetic markers associate with atrial fibrillation risk in Europeans and Japanese. J Am Coll Cardiol. 2014 Apr 1;63(12):1200–10.

21. Wang J, Klysik E, Sood S, Johnson RL, Wehrens XH, Martin JF. Pitx2 prevents susceptibility to atrial arrhythmias by inhibiting left-sided pacemaker specification. Proc Natl Acad Sci U S A. 2010; 107: 9753–9758.

22. Kirchhof P, Kahr PC, Kaese S, Piccini I, Vokshi I, Scheld HH, Rotering H, Fortmueller L, Laakmann S, Verheule S, Schotten U, Fabritz L, Brown NA. PITX2c is Expressed in the Adult Left Atrium, and Reducing Pitx2c Expression Promotes Atrial Fibrillation Inducibility and Complex Changes in Gene Expression. Circ Cardiovasc Genet. 2011;4:123–33.

23. Chinchilla A, Daimi H, Lozano-Velasco E, Dominguez JN, Caballero R, Delpon E, Tamargo J, Cinca J, Hove-Madsen L, Aranega AE, Franco D. Pitx2 insufficiency leads to atrial electrical and structural remodelling linked to arrhythmogenesis. Circ Cardiovasc Genet. 2011;4:269–79.

24. Tao Y, Zhang M, Li L, Bai Y, Zhou Y, Moon AM, Kaminski HJ, Martin JF. Pitx2, an atrial fibrillation predisposition gene, directly regulates ion transport and intercalated disc genes. Circ Cardiovasc Genet. 2014 Feb;7(1):23–32

25. Lozano-Velasco E, Hernández-Torres F, Daimi H, Serra SA, Herraiz A, Hove-Madsen L, Aránega A, Franco D. Pitx2 impairs calcium handling in a dose-dependent manner by modulating Wnt signalling. Cardiovasc Res. 2016; 109(1):55–66.

26. Torrado M, Franco D, Hernández-Torres F, Crespo-Leiro MG, Iglesias-Gil C, Castro-Beiras A, Mikhailov AT. Pitx2c is reactivated in the failing myocardium and stimulates myf5 expression in cultured cardiomyocytes. PLoS One. 2014 Mar 4;9(3):e90561. doi:10.1371/journal.pone.0090561.

27. Chen J, Kubalak SW and Chien KR. Ventricular muscle-restricted targeting of the RXRalpha gene reveals a non-cell-autonomous requirement in cardiac chamber morphogenesis. Development. 1998;25:1943–1949.

28. Livak KJ, Schmittgen TD. Analysis of relative gene expression data using real-time quantitative PCR and the 2(-Delta Delta C(T)) Method Methods. 2001;25:402–408.

29. Domínguez JN, Navarro F, Franco D, Thompson RP, Aránega AE. Temporal and spatial expression pattern of beta1 sodium channel subunit during heart development. Cardiovasc Res. 2005;65:842–850.

30. Goldbarg AN, Hellerstein HK, Bruell JH, Daroczy AF. Electrocardiogram of the normal mouse, Mus musculus: general considerations and genetic aspects. Cardiovasc Res.. 1968; 2(1):93–99.

31. Salama G, London B. Mouse models of long QT syndrome. J Physiol. 2007;578:43–53.

32. Speerschneider T, Thomsen MB. Physiology and analysis of the electrocardiographic T wave in mice. Acta Physiol.. 2013; 209(4):262–71.

33. Boukens BJ, Rivaud MR, Rentschler S, Coronel R. Misinterpretation of the mouse ECG: ‘musing the waves of Mus musculus’. J Physiol. 2014;592:4613–26.

34. Furtado MB, Biben C, Shiratori H, Hamada H, Harvey RP. Characterization of Pitx2c expression in the mouse heart using a reporter transgene. Dev Dyn.. 2011; 240(1):195–203.

35. Hernandez Torres F, Franco D, Aranega AE, Navarro F. Expression patterns and immunohistofluorescence localization of Pitx2b transcription factor in the developing mouse heart. Int J Dev Biol. 2015;59(4-6):247–54.

36. Johnson JN, Grifoni C, Bos JM, Saber-Ayad M, Ommen SR, Nistri S, Cecchi F, Olivotto I, Ackerman MJ. Prevalence and clinical correlates of QT prolongation in patients with hypertrophic cardiomyopathy. Eur Heart J. 2011;32:1114–1120.

37. Kang YJ. Cardiac hypertrophy: a risk factor for QT-prolongation and cardiac sudden death. Toxicol Pathol.. 2006; 34(1):58–66.

38. Nerbonne JM. Studying cardiac arrhythmias in the mouse‐‐a reasonable model for probing mechanisms? Trends Cardiovasc Med. 2004;14:83–93.

39. Rose J, Armoundas AA, Tian Y, DiSilvestre D, Burysek M, Halperin V, O’Rourke B, Kass DA, Marbán E, Tomaselli GF. Molecular correlates of altered expression of potassium currents in failing rabbit myocardium. Am J Physiol Heart Circ Physiol. 2005; 288:H2077–87.

40. Bignolais O, Quang KL, Naud P, El Harchi A, Briec F, Piron J, Bourge A, Leoni AL, Charpentier F, Demolombe S. Early ion-channel remodeling and arrhythmias precede hypertrophy in a mouse model of complete atrioventricular block. J Mol Cell Cardiol. 2011;51:713–721.

41. Liao X, Haldar SM, Lu Y, Jeyaraj D, Paruchuri K, Nahori M, Cui Y, Kaestner KH, Jain MK. Krüppel-like factor 4 regulates pressure-induced cardiac hypertrophy. J Mol Cell Cardiol. 2010 Aug;49(2):334–8.

42. Xie C, Huang H, Sun X, Guo Y, Hamblin M, Ritchie RP, Garcia-Barrio MT, Zhang J, Chen YE. MicroRNA-1 regulates smooth muscle cell differentiation by repressing Kruppel-like factor 4. Stem Cells Dev. 2011 Feb;20(2):205–10.

43. de Lange FJ, Moorman AFM, Christoffels, VM. Atrial cardiomyocyte-specific expression of Cre recombinase driven by an Nppa gene fragment. Genesis. 2003;37:1–4.

44. Martínez-Fernandez S, Hernández-Torres F, Franco D, Lyons GE, Navarro F, Aránega AE. Pitx2c overexpression promotes cell proliferation and arrests differentiation in myoblasts. Dev Dyn. 2006;235:2930–2939.

45. Gherzi R, Trabucchi M, Ponassi M, Gallouzi I-E, Rosenfeld MG, Briata P. Akt2-mediated phosphorylation of Pitx2 controls Ccnd1 mRNA decay during muscle cell differentiation. Cell Death Differentiation. 2010;17:975–983.

46. Kioussi C, Briata P, Baek SH, Rose DW, Hamblet NS, Herman T, Ohgi KA, Lin C, Gleiberman A, Wang J, Brault V, Ruiz-Lozano P, Nguyen HD, Kemler R, Glass CK, Wynshaw-Boris A, Rosenfeld MG. Identification of a Wnt/Dvl/beta-Catenin ‐‐ > Pitx2 pathway mediating cell-type-specific proliferation during development. Cell. 2002;111: 673–685.

47. Lozano-Velasco E, Chinchilla A, Martínez-Fernández S, Hernández-Torres F, Navarro F, Lyons GE, Franco D, Aránega AE. Pitx2c modulates the cardiac trancriptional machinery in differentiating cardiomyocytes from murine embryonic stem cells. Cell Tissue Organs. 2011; 194(5):349–62.

48. Galli D, Domínguez JN, Zaffran S, Munk A, Brown NA, Buckingham ME. Atrial myocardium derives from the posterior region of the second heart field, which acquires left-right identity as Pitx2c is expressed. Development. 2008;135:1157–1167.

49. Shih HP, Gross MK, Kioussi C. Cranial muscle defects of Pitx2 mutants result from specification defects in the first branchial arch. Proc Natl Acad Sci U S A. 2007;104: 5907–5912.

50. Pérez-Hernández M, Matamoros M, Barana A, Amorós I, Gómez R, Núñez M, Sacristán S, Pinto Á, Fernández-Avilés F, Tamargo J, Delpón E, Caballero R. Pitx2c increases in atrial myocytes from chronic atrial fibrillation patients enhancing IKs and decreasing ICa,L. Cardiovasc Res. 2016 Mar 1;109(3):431–41.

51. Nishizaki M, Hiraoka M. Gene mutations associated with atrioventricular block complicated by long QT syndrome. Circ J. 2010;74:2546–2547.

52. Furushima H, Chinushi M, Sato A, Aizawa Y, Kikuchi A, Takakuwa K, Tanaka K. Fetal atrioventricular block and postpartum augmentative QT prolongation in a patient with long-QT syndrome with KCNQ1 mutation. J Cardiovasc Electrophysiol. 2010;21:1170–1173.

53. Demolombe S, Lande G, Charpentier F, van Roon MA, van den Hoff MJ, Toumaniantz G, et al. Transgenic mice overexpressing human KvLQT1 dominant-negative isoform. Part I: Phenotypic characterisation. Cardiovasc Res. 2001;50:314–327.

54. Oka Y, Itoh H, Ding WG, Shimizu W, Makiyama T, Ohno S, Nishio Y, Sakaguchi T, Miyamoto A, Kawamura M, Matsuura H, Horie M. Atrioventricular block-induced Torsades de Pointes with clinical and molecular backgrounds similar to congenital long QT syndrome. Circ J. 2010;74: 2562–2571.

55. Lozano-Velasco E, Wangensteen R, Quesada A, Garcia-Padilla C, Osorio JA, Ruiz-Torres MD, Aranega A, Franco D. Hyperthyroidism, but not hypertension, impairs PITX2 expression leading to Wnt-microRNA-ion channel remodeling. PLoS One. 2017 Dec 1;12(12):e0188473. doi:10.1371/journal.pone.0188473.

56. Zhao Y, Ransom JF, Li A, Vedantham V, von Drehle M, Muth AN, Tsuchihashi T, McManus MT, Schwartz RJ, Srivastava D. Dysregulation of cardiogenesis, cardiac conduction, and cell cycle in mice lacking miRNA-1-2. Cell. 2007;129:303–317.

57. Yang B, Lin H, Xiao J, Lu Y, Luo X, Li B, Zhang Y, Xu C, Bai Y, Wang H, Chen G, Wang Z. The muscle-specific microRNA miR-1 regulates cardiac arrhythmogenic potential by targeting GJA1 and KCNJ2. Nat Med. 2007;13:486–491.

58. Girmatsion Z, Biliczki P, Bonauer A, Wimmer-Greinecker G, Scherer M, Moritz A, Bukowska A,Goette A, Nattel S, Hohnloser SH, Ehrlich JR. Changes in microRNA-1 expression and IK1 up-regulation in human atrial fibrillation. Heart Rhythm. 2009;6:1802–1809.

59. Lu Y, Zhang Y, Wang N, Pan Z, Gao X, Zhang F, Zhang Y, Shan H, Luo X, Bai Y, Sun L, Song W, Xu C, Wang Z, Yang B. MicroRNA-328 contributes to adverse electrical remodeling in atrial fibrillation. Circulation. 2010 Dec 7;122(23):2378–87.

